# UV radiation increases flavonoid protection but decreased reproduction in *Silene littorea*

**DOI:** 10.1101/2020.03.30.016295

**Authors:** J. C. Del Valle, M. L. Buide, J. B. Whittall, F. Valladares, E. Narbona

## Abstract

Plants respond to changes in ultraviolet (UV) radiation via morphological and physiological changes. Among the variety of plant UV-responses, the synthesis of UV-absorbing flavonoids constitutes an effective non-enzymatic mechanism to mitigate photoinhibitory and photooxidative damage caused by UV stress, either reducing the penetration of incident UV radiation or acting as quenchers of reactive oxygen species (ROS). In this study, we designed a UV-exclusion experiment to investigate the effects of UV radiation in *Silene littorea*. We spectrophotometrically quantified concentrations of both anthocyanins and non-anthocyanin flavonoids (flavones) in petals, calyces, leaves and stems. Furthermore, we analyzed the UV effect on the photosynthetic activity in hours of maximum solar radiation and we tested the impact of UV radiation on male and female reproductive performance. We found that anthocyanin concentrations showed a significant decrease of about 20% with UV-exclusion in petals and stems, and 30% in calyces. Flavone concentrations showed a significant reduction of approximately 25% in calyces and stems, and 12% in leaves. Photochemical efficiency of plants grown under UV stress decreased sharply at maximum light stress, but their ability for recovery after light-stress was not affected. In addition, exposure to UV radiation does not seem to affect ovule production or seed set, but decreases the total seed production per plant and pollen production by 69% and 31%, respectively. Our results demonstrate that UV radiation produced opposite effects on flavonoid accumulation and reproduction in *S. littorea*. UV stress increased flavonoid concentrations, suggesting a photoprotective role of flavonoids against UV radiation, but had negative consequences for reproduction. We propose that this trade-off helps this species to occupy exposed habitats with high UV radiation.

## Introduction

Ultraviolet (UV) radiation can both help and harm plants. Many flowering plants rely on UV nectar guides for pollination services [1]. Simultaneously, the high energy of UV radiation can be damaging to cells and presents a unique abiotic challenge to most land plants [2]. Furthermore, the “invisible” nature of UV radiation makes it particularly enigmatic at the ecological and physiological scales. UV-A (315 – 400 nm) and UV-B (280 – 315 nm) radiation has numerous positive and negative effects at the cellular and organismal scales [3–6], inducing a variety of morphological responses in plants [4,7]. In addition, UV-B radiation exerts damaging effects on DNA and chloroplasts, particularly photosystem II (PSII), and indirectly generate reactive oxygen species (ROS) that can further damage the photosynthetic apparatus [8,9].

Plants have developed a variety of mechanisms to avoid the harmful effects of UV radiation, mainly through the screening of UV wavelengths, the repairing of UV-induced damage or the quenching of ROS [6,8,9]. Although the latter is primarily performed by antioxidant enzymes that control ROS levels [8,10], flavonoids and other phenolic compounds are involved in the detoxification of ROS as well [11–13]. Flavonoids are potent scavengers of ROS that prevent lipid peroxidation and scavenge free radicals, especially those flavonoids having a catechol group in the B-ring of the flavonoid skeleton (e.g. quercetin derivatives) [14,15]. Furthermore, exposure to excess light or UV-B radiation increases the synthesis of effective antioxidant dihydroxy B-ring-substituted flavonoids (e.g. luteolin derivatives) at expense of other less effective antioxidant flavonoids (e.g. kaempferol derivatives) [16,17]. In addition to their key antioxidant functions, other studies have attributed to flavonoids an important role in photoprotection through the screening of the UV radiation [18,19]. For example, epidermal flavonols play a predominant role in UV-B screening in leaves of *Secale cereale* and *Centella asiatica* [20,21].

Within the flavonoids, anthocyanins are plant pigments that are synthesized in the last steps of the flavonoid biosynthetic pathway [22]. Anthocyanins mainly absorb in the green region of visible (VIS) spectrum (500-565 nm), reducing the overall photosynthetically active radiation (PAR) (400 - 700nm) hitting the chloroplasts and helping plants to have a faster photosynthetic recovery after saturating light stress [23–25]. In addition, when anthocyanins are acylated, they can absorb UV radiation, and may contribute to ROS scavenging even more than other phenolic compounds [26–29]. Yet, UV stress is known to induce anthocyanin biosynthesis, which may contribute to the tolerance to UV radiation [30,31].

The aforementioned photoprotective functions of UV-induced flavonoids are not restricted to photosynthetic tissues, but also occur in floral structures such as anthers, ovaries, petals and sepals. Pollen grains accumulate flavonoids to protect them from UV-B damage and preserve their viability after anthesis [32], whereas flavonoids protect ovules by shielding ovaries from UV radiation [33]. In the same way, the accumulation of protective flavonoids in petals and sepals can reduce the damaging effects of UV radiation on these and other nearby reproductive tissues [34]. Additionally, petal flavonoids form a UV pigmentation floral pattern to guide pollinators to nectaries, thus UV-induced changes in the size of nectar guides might affect the pollination activity [1,35]. In addition, UV radiation may induce a variety of plant morphological responses in these reproductive structures. Many studies have reported a complex UV dose-response involving differential effects on reproductive morphological traits (reviewed in [7]). For example, Koti et al. reported that UV-B radiation negatively affected the flower phenology, pollen germination and its viability in soybean (*Glycine max*) [36], and similarly decreased pollen and flower production over time in *Brassica rapa* [37].

This paper describes the effects of UV-radiation in flavonoid accumulation and reproduction of the shore campion (*Silene littorea* Brot., Caryophyllaceae). This annual species is endemic to coastal foredunes along the Iberian Peninsula and accumulates flavonoids (flavones and anthocyanins) in petals, calyces, stems and leaves [38,39]. Our previous work has shown a latitudinal gradient in flavonoid accumulation that tends to increase from north to south in most plant tissues - correlated with increased solar exposure and temperatures [39]. Moreover, we found that intense solar radiation, including UV and VIS spectra, increased the synthesis of flavones and anthocyanins in most aboveground tissues of *S. littorea* [40]. In this study, we focus on the effect of the UV irradiation on flavonoid accumulation in this species. We quantified the concentrations of flavones and anthocyanins in petals, calyces, leaves and stems of plants grown with or without exposure to UV radiation. Then, we analyzed the UV effects on the photosynthetic efficiency and the male and female reproductive output.

Flavonoids have a key role in photoprotection [15,19], but the synthesis of these compounds may represent a cost for the plant [24]. Consequently, we predict that the exclusion of UV radiation will result in a decrease in UV-inducible flavonoid concentrations in all tissues. This energetic and carbon savings under UV-exclusion may result in increased reproductive allocation [41]. In contrast, without UV protection, we predict that photodamage will decrease photosynthetic activity [9,42] and show lower reproductive output. Since *S. littorea* inhabits exposed coastal dunes habitats with high solar radiation levels, we hypothesize that this species will have effective light-stress recovery system that prevents long-term photoinhibition.

## Materials and methods

### Study system and experimental design

*Silene littorea* is an annual plant that accumulates anthocyanins (cyanidin derivatives) and flavones (mainly isovitexin and isoorientin derivatives) in both reproductive and vegetative tissues [38] (Fig 1). This species inhabits coastal populations from the northwestern corner to the southeastern portion of the Iberian Peninsula [39]. We collected seeds of six plants from a northwestern population (Furnas; 42° 38’ 15’’ N, 9° 2’ 21’’ W) and six plants from a southwestern population (Sines; 37° 55’ 17’’ N, 8° 48’ 17’’ W). These two populations, which are separated by approximately 500 km along the Atlantic coast of the Iberian Peninsula, are exposed to a different degree of solar irradiance, being approximately 30% higher in southern latitudes [39].

**Fig 1.**
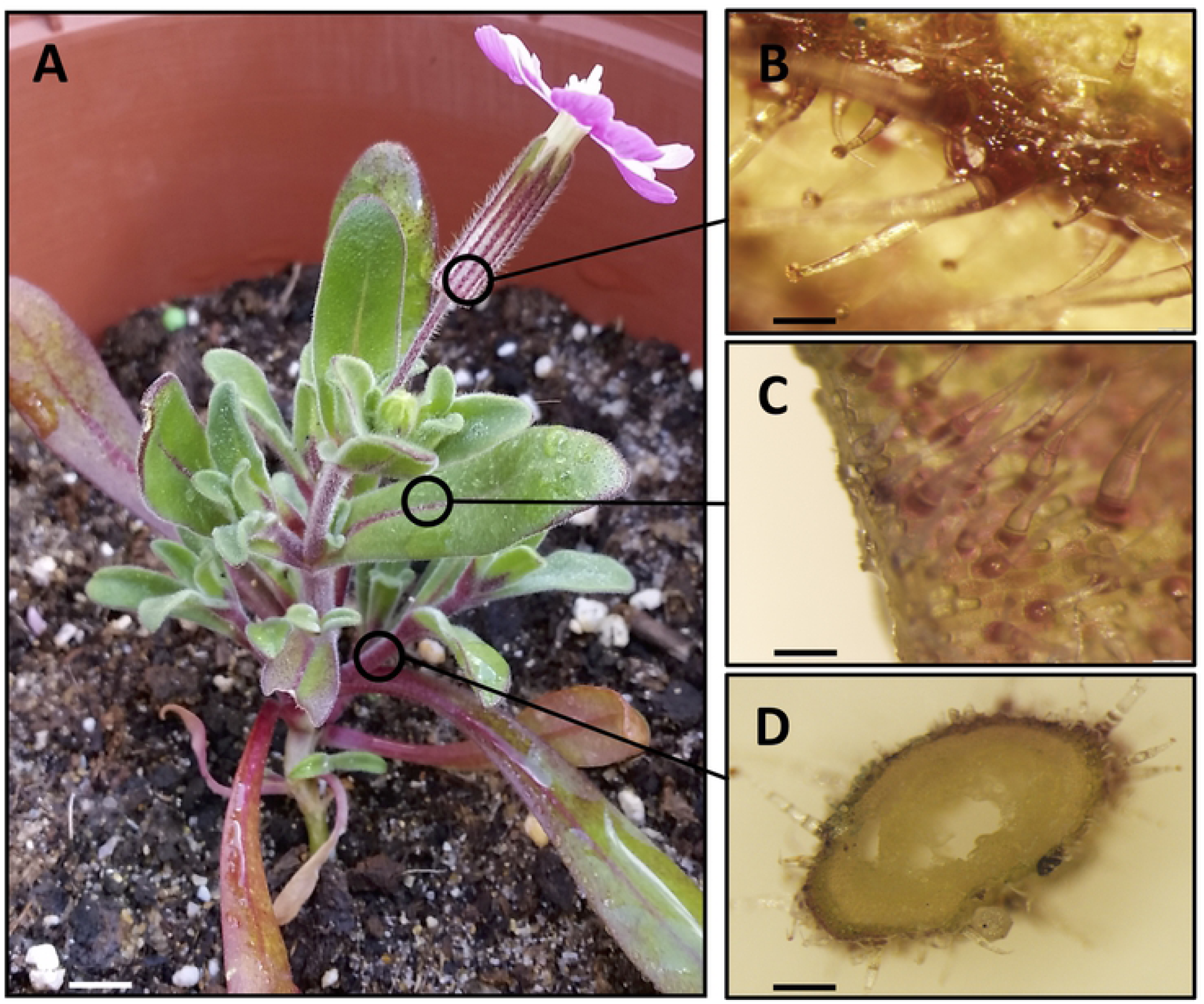
Details of a *Silene littorea* plant (A) to show the accumulation of anthocyanins throughout the whole plant. B, C and D showed stereo-microscope photographs of surface of the calyx ribs, adaxial surface of the leaf, and cross section of a stem, respectively. Scale bar: 5 mm (A), 0.5 mm (B, C), and 1 mm (D).

Seeds obtained from the 12 maternal families were scarified and maintained at 45 °C for a month to break dormancy, and afterwards they were germinated in a germination chamber at 22 °C/15 °C (12 h light/12 h dark). The resulting seedlings were planted in pots filled with 2.5 L of a mixture of standard substrate (80-90% organic material, pH = 6.5) and beach sand (v:v 50:50) and were grown in the greenhouse at Pablo de Olavide University (Seville, Spain) for one month. In February 2016 (one month before flowering), pots were put outside on two benches in the experimental garden. The bench assigned to the UV-present treatment was covered with a methacrylate filter that transmitted 100% UV irradiance, whereas the bench assigned to the UV-exclusion treatment was covered with a polycarbonate filter preventing most UV radiation (< ∼ 385nm). The UV-exclusion treatment produced a reduction of 9.2% and 100% of total transmitted sunlight and UV radiation, respectively. Maximum solar irradiation at natural sunlight was 1258 W/m^2^ and UV irradiance was 4.36 W/m^2^. Measures were taken at 14:00 h of a sunny day (6^th^ June 2016). Total solar radiance and UV were measured by means of Megger PVM210 irradiance meter (range sensitivity = 1999 W/m^2^; resolution = 0.1 W/m^2^) (Megger Co., Dallas, USA) and PCE-UV34 UV light meter (range sensitivity = 0.000 to 19.9 W/m^2^; resolution = 0.01 W/m^2^) (PCE Inst., Durham, UK), respectively. Given the low germination rates of this species and the high mortality at the seedling stage, the final number of surviving plants was 65 (belonging to nine maternal families). These plants were assigned to the UV-present (41) and UV-exclusion (24) treatments, respectively (S1 Table). Plants that shared the same maternal family were equally assigned to each treatment whenever possible.

### Flavonoid quantification

During peak flowering (May 2016), samples of petals, calyces (five petals and the calyx of the same flower), leaves (selected mid-stem) and stems (1 cm length section from the main axis) were collected from 34 and 22 plants grown in the UV-present and UV-exclusion environments, respectively (S1 Table). Samples were extracted in 1.5 ml of MeOH containing 1% of HCl following the procedure described in Del Valle et al. [39]. Three replicates of 200 μL per sample extraction were used to estimate flavonoid concentrations using a Multiskan GO microplate spectrophotometer (Thermo Fisher Scientific Inc., MA, USA). Anthocyanins and flavones were quantified at A_520_ and A_350_, respectively. In photosynthetic organs (calyces, leaves and stems), anthocyanin concentration was corrected as A_520_ - (0.24 x A_653_) to compensate for the small overlap absorption by chlorophyll [43]. Anthocyanins and flavones concentrations were calculated following Del Valle et al. [38] and expressed as milligrams of cyanidin-3-glucoside, isovitexin and isoorientin equivalents per gram fresh weight, respectively.

### Assessment of photosynthetic activity

To determine if there were physiological consequences of plants grown with and without UV radiation, the photochemical efficiency of PSII (*Fv/Fm*) was measured in calyces and leaves of 30 plants from Sines (14 and 16 from the UV-present and UV-exclusion treatments, respectively) using a field portable pulse-modulated chlorophyll fluorometer (FMS2, Hansatech Instruments, Norfolk, UK). Measurements were carried out in predawn (∼ 0700h) and in maximum solar radiation (∼1430h), and in the early (March) and maximum (May) stages of the flowering period. To asses the physiological status of photosynthetic tissues across the experiment, measurements were carried out in fully exposed plants on two sunny days [44]. To minimize temporal variation in *Fv/Fm*, all measurements were made in a period of one hour. Prior to taking physiological measurements, samples were acclimated for 30 minutes in dark using leaf-clips that contain a mobile shutter.

### Assessment of plant reproductive performance

Flower and fruit production in 41 and 24 plants from the UV-present and UV-exclusion treatments were monitored weekly during the entire flowering period, from March 10^th^ to June 20^th^. These individual flowers were monitored for either fruit production or fruit abortion to determine the proportion of flowers that set fruit. In May, one mature fruit per plant was collected if possible. A total of 33 and 21 mature fruits were collected from plants growing in the UV-present and UV-exclusion treatments, respectively. For each mature fruit, their seeds and aborted ovules were counted under the dissecting microscope to calculate the proportion of ovules that set seed. Then, seed production per plant was estimated for those plants from which we collected mature fruits and calculated as the product of seeds/fruit x total number of fruits produced during the flowering period. Pollen and ovule production were analyzed following the procedure described in Narbona et al. [45] from unopened flower buds preserved in FAA (95 % ethanol, dH_2_O, 37-40 % formaldehyde, acetic glacial acid, 10:7:2:1) of nine and 13 plants grown in the UV-present and UV-exclusion treatments, respectively. The total number of pollen grains per anther was calculated as the average of pollen grains counted in one upper and one lower anther of an unopened flower bud per plant.

### Statistical analysis

Generalized linear mixed models (GLMMs) with Gaussian link functions were used to test the effect of UV radiation on the accumulation of anthocyanins and flavones in each plant tissue, considering treatment and population as fixed factors and maternal family as a random factor. Flavonoid concentrations were log-transformed prior to conduct the GLMMs analysis. Pairwise comparisons between UV-present and UV-exclusion treatments were carried out using the “multcomp” R-package and its “cld” (compact letter display) function was used to show differences between populations [46]. Due to the low number of experimental plants, we used the conservative Bonferroni adjustment of p-values in pairwise comparisons [47]. The same analyses were used to test for differences in male and female reproductive performance and in the photochemical efficiency of PSII (*Fv/Fm*) between plants grown in the different UV treatments. For this latter analysis, independent comparisons were done for leaves and calyces and in the early (March) and maximum (May) stages of the experiment, as well as pairwise comparisons of the photochemical efficiency between predawn and afternoon conditions. Pearson’s correlations with a Bonferroni adjustment for multiple comparisons were used to assess the relationship between flavonoid production and male and female reproductive output [48]. All analyses were performed in R v3.4.0 [49]. GLMMs were carried out using the R library “lme4” [50].

## Results

### Effects of UV radiation on flavonoid production

In general, plants from the UV-exclusion treatment showed lower accumulation of anthocyanins, but this decrease was not homogenous in all tissues. Specifically, anthocyanin concentrations in petals and stems decreased approximately 20% in these plants, whereas in calyces the decrease was of 30% and the differences were marginally significant (Fig 2, Table 1). Anthocyanins were nearly absent altogether in leaves (Fig 2E). Flavone concentrations in plants from the UV-exclusion treatment were lower by 12%, 23%, and 25% in leaves, calyces, and stems, respectively, but in petals the differences were not significant (Table 1).

**Table 1.** Results from GLMMs testing the effect of UV radiation, population and their interaction on the production of anthocyanins and flavones in each plant tissue.

|  |  | Anthocyanins |  |  |  |  | Flavones |  |  |  |  |
| --- | --- | --- | --- | --- | --- | --- | --- | --- | --- | --- | --- |
| Tissue | Source of variation | SS | Numerator d.f. | Denominator d.f. | F | P | SS | Numerator d.f. | Denominator d.f. | F | P |
| Petals | Treatment | 0.670 | 1 | 46.62 | 7.968 | **0.007** | 0.019 | 1 | 38.78 | 1.469 | 0.233 |
|  | Population | 0.033 | 1 | 19.79 | 0.396 | 0.536 | 0.477 | 1 | 42.86 | 36.15 | **<0.001** |
|  | Treatm. x Pop. | 0.096 | 1 | 47.86 | 1.140 | 0.291 | 0.001 | 1 | 38.68 | 0.004 | 0.949 |
| Calyces | Treatment | 0.674 | 1 | 44.48 | 3.729 | 0.059 | 1.013 | 1 | 44.39 | 37.27 | **<0.001** |
|  | Population | 0.121 | 1 | 43.22 | 0.667 | 0.419 | 0.606 | 1 | 46.06 | 22.29 | **<0.001** |
|  | Treatm. x Pop. | 0.231 | 1 | 45.11 | 1.276 | 0.265 | 0.018 | 1 | 44.94 | 0.647 | 0.426 |
| Leaves | Treatment | 0.007 | 1 | 46.70 | 0.176 | 0.677 | 0.368 | 1 | 49.00 | 5.145 | **0.028** |
|  | Population | 0.078 | 1 | 23.18 | 1.899 | 0.181 | 1.166 | 1 | 49.00 | 16.28 | **<0.001** |
|  | Treatm. x Pop. | 0.199 | 1 | 48.29 | 4.840 | **0.033** | 0.023 | 1 | 49.00 | 0.327 | 0.570 |
| Stems | Treatment | 1.572 | 1 | 44.65 | 8.527 | **0.005** | 1.295 | 1 | 46.10 | 6.203 | **0.016** |
|  | Population | 0.035 | 1 | 49.27 | 0.191 | 0.664 | 0.049 | 1 | 42.77 | 0.237 | 0.629 |
|  | Treatm. x Pop. | 0.339 | 1 | 45.22 | 1.837 | 0.182 | 0.008 | 1 | 46.92 | 0.041 | 0.841 |

Significant *P*-values were highlighted in bold.

**Fig 2.**
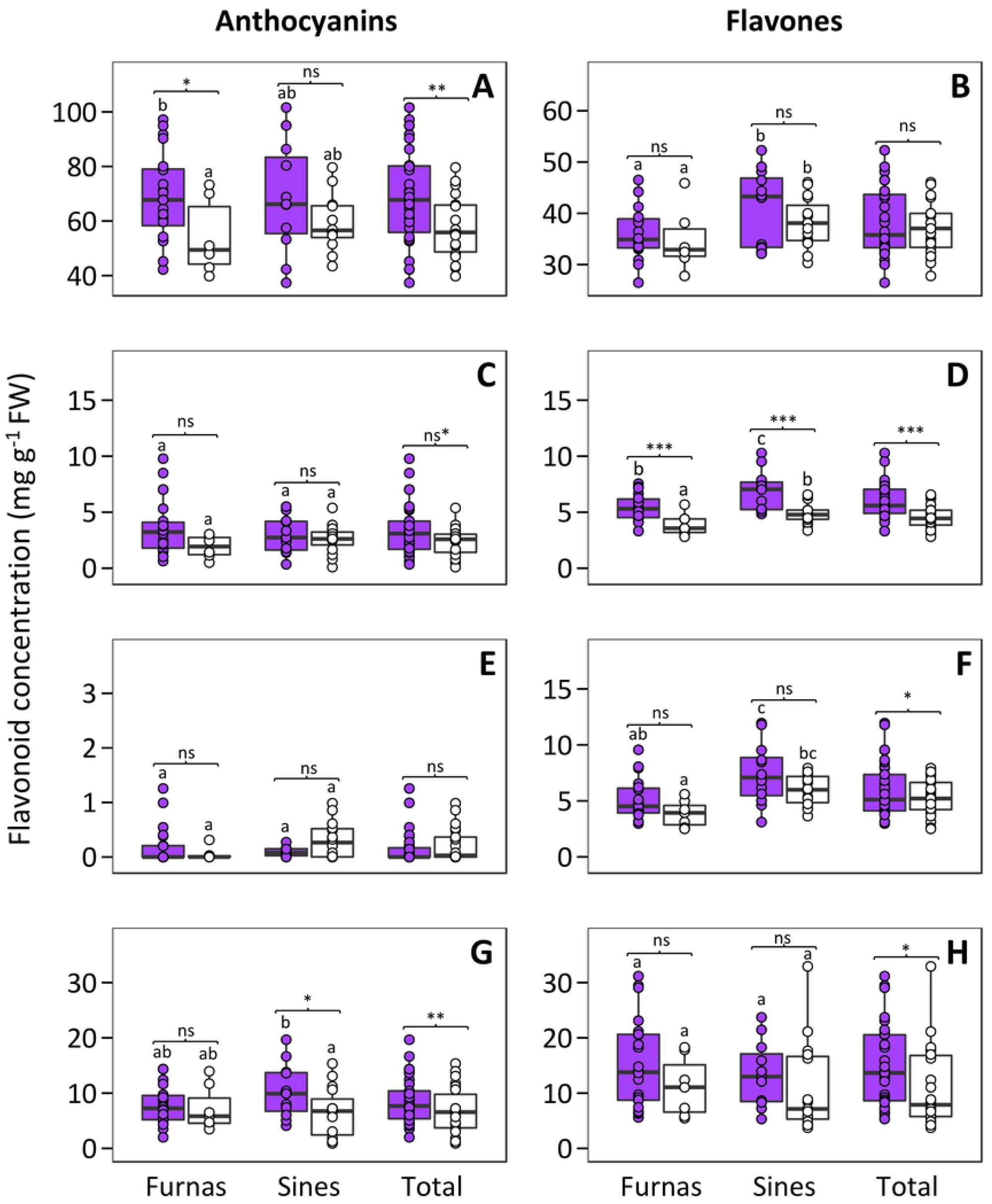
Boxplots representing anthocyanin and flavone concentrations in the UV-present (purple boxes) and UV-exclusion (white boxes) treatments in petals (A, B), calyces (C, D), leaves (E, F) and stems (G, H). The central line displays the median, the bottom and top of the box are the first and third quartiles, and dots represent sample values. Lowercase letters are used to show results of multiple comparisons between populations. Within each population, pairwise comparisons between light treatments using Bonferroni adjustment are showed. FW, fresh weight; ns, not significant; ns*, marginally significant; *, *P* < 0.05; **, *P* < 0.01; ***, *P* < 0.001.

Sines and Furnas populations did not show significant differences in anthocyanin concentrations in any of the sampled tissues (Table 1). Conversely, the flavone concentrations were significantly higher in plants from Sines, the southern population, in all tissues except for the stems (Fig 2, Table 1), and the interactions of light treatment and population were not significant (i.e. the decrease of flavone concentration in UV-exclusion plants was homogeneous in both populations).

When we analyzed each population independently, we found that the only significant differences in anthocyanin concentrations between treatments were in petals of plants from Furnas and in stems of plants from Sines (Figs 2A and G). With respect to flavones concentrations, the only significant differences between treatments were found in calyces of plants from both populations (Fig 2D). Interestingly, plants from both UV-exclusion and UV-present treatments of Sines showed higher levels of flavones than their respective treatments in Furnas.

### Effects of UV radiation in photosynthetic performance

Plants decreased their photochemical efficiency (*Fv/Fm*) in the afternoon, when plants were maximally exposed to light stress, but in the predawn, after a whole night of relaxation of photoinactivation, they showed *Fv/Fm* values within the range of healthy plants (∼ 0.85) (Fig 3, Table 2). Leaves showed significant differences in their photochemical efficiency between UV-treatments and between measurement conditions (predawn or afternoon), and the interaction of UV-treatments and measurement conditions was also significant (Table 2). In the afternoon, leaves of the UV-exclusion treatment showed a 20.8% and 57.4% reduction of *Fv/Fm* values in early (March) and maximum (May) stages of the flowering period, respectively (*P* < 0.001 for both pairwise comparisons, Table 3; Fig 3A and B). In calyces, statistical differences in their photochemical efficiency were found only between measurement conditions (predawn or afternoon) both in March and May (Table 2). Pairwise comparisons in calyces revealed significant lower *Fv/Fm* values in afternoon conditions, regardless of the UV treatment or the flowering period (*P* < 0.032, Table 3; Fig 3C and D).

**Table 2.** Results from GLMMs testing the effect of UV radiation, measurement condition (predawn or afternoon) and their interaction on the photochemical efficiency of PSII (*Fv/Fm*) in leaves and calyces.

| Tissue | Stage | Source of variation | SS | Numerator d.f. | Denominator d.f. | F | P |
| --- | --- | --- | --- | --- | --- | --- | --- |
| Leaves | Early (March) | Treatment | 0.030 | 1 | 50.00 | 19.07 | < 0.001 |
|  |  | Measurement condition | 0.056 | 1 | 50.00 | 34.86 | < 0.001 |
|  |  | Treatm. x Measurement condition | 0.020 | 1 | 50.00 | 12.34 | < 0.001 |
|  | Maximum (May) | Treatment | 0.083 | 1 | 40.00 | 6.209 | 0.017 |
|  |  | Measurement condition | 0.528 | 1 | 40.00 | 39.50 | < 0.001 |
|  |  | Treatm. x Measurement condition | 0.115 | 1 | 40.00 | 8.614 | 0.006 |
| Calyces | Early<br>(March) | Treatment | 0.004 | 1 | 32.39 | 2.822 | 0.103 |
|  |  | Measurement condition | 0.039 | 1 | 47.25 | 30.42 | **< 0.001** |
|  |  | Treatm. x Measurement condition | 0.004 | 1 | 47.25 | 3.401 | 0.071 |
|  | Maximum<br>(May) | Treatment | 0.001 | 1 | 40.39 | 0.182 | 0.672 |
|  |  | Measurement condition | 0.375 | 1 | 38.80 | 48.32 | **< 0.001** |
|  |  | Treatm. x Measurement condition | 0.005 | 1 | 38.80 | 0.639 | 0.429 |

Significant *P*-values were highlighted in bold.

**Table 3.** Comparisons of the photochemical efficiency (*Fv/Fm*) between predawn and afternoon conditions. Pairwise comparisons were independently performed in leaves and calyces from the UV-exclusion and UV-present treatments and either in the early (March) and maximum (May) stages of flowering period.

Significant *P*-values were highlighted in bold.
|  |  |  |  |  |  |  |
| --- | --- | --- | --- | --- | --- | --- |
| Tissue | Stage | Treatment | Estimate | Std. Error | Z value | *P* |
| Leaves | Early (March) | UV-exclusion | -0.027 | 0.014 | -1.873 | 0.239 |
|  |  | UV-present | -0.104 | 0.017 | -6.117 | **< 0.001** |
|  | Maximum (May) | UV-exclusion | -0.117 | 0.047 | -2.480 | 0.063 |
|  |  | UV-present | -0.323 | 0.052 | -6.252 | **< 0.001** |
| Calyces | Early (March) | UV-exclusion | -0.037 | 0.013 | -2.788 | **0.032** |
|  |  | UV-present | -0.073 | 0.015 | -4.889 | **< 0.001** |
|  | Maximum (May) | UV-exclusion | -0.163 | 0.035 | -4.611 | **< 0.001** |
|  |  | UV-present | -0.205 | 0.039 | -5.202 | **< 0.001** |

**Fig 3.**
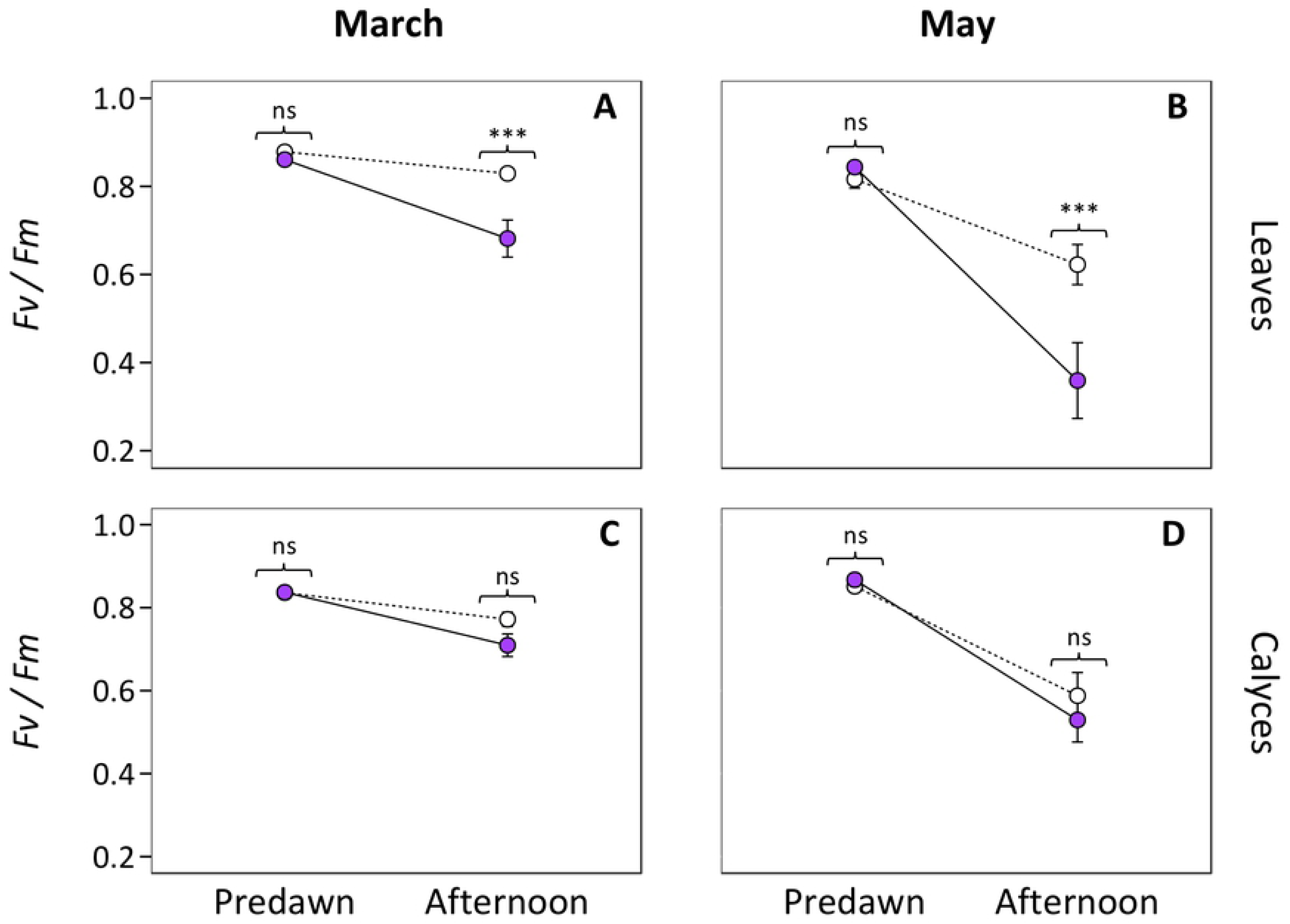
Variation of photochemical efficiency (*Fv*/*Fm*) from predawn conditions to afternoon. The mean *Fv/Fm* values obtained from leaves (A, B) and calyces (C, D) in the early (March; A and C) and maximum (May; B and D) stages of the flowering period are showed. Plants from the UV-present treatment are displayed by pink filled circles and solid lines, whereas those from the UV-exclusion treatment are displayed with empty circles and dashed lines. Results of independent pairwise comparisons after Bonferroni corrections between UV treatments in predawn and afternoon conditions are displayed. Error bars represent ± SE.

### Effects of UV radiation in reproductive performance

Flower production showed statistical differences between the two experimental conditions (Table 4). Plants from the UV-exclusion treatment displayed approximately five times more flowers than those with UV-present (261.4 ± 30.1 and 50.7 ± 8.3, respectively; mean ± SE; Fig 4A). In addition, flower production was significantly different for both populations, being higher in Sines plants. Conversely, fruit set was statistically higher in the UV-present treatment and in plants from Furnas population (Fig 4B, Table 4). The number of ovules per flower and seed set was statistically similar between light treatments or populations (Figs 4C and D). The total seed production per plant was approximately three times higher in plants from the UV-exclusion treatment compared to the UV-present plants (46.2 ± 7.8 and 14.3 ± 1.9, respectively; Fig 4E), and did not show statistical differences between populations (Table 4). Pollen production decreased by ∼31% in plants exposed to UV radiation (2126.1 ± 99.0 and 1473.9 ± 85.8, respectively; Fig 4F), but again differences between populations was not found. The interactions of UV treatment and population were not significant for any of the studied reproductive outputs (Table 4).

**Table 4.** Results from GLMMs testing the effect of UV radiation, population and their interaction on the estimations of male and female reproductive performance in *S. littorea*.

|  | Source of variation | SS | Numerator d.f. | Denominator d.f. | F | P |
| --- | --- | --- | --- | --- | --- | --- |
| Flowers per plant | Treatment | 37.55 | 1 | 52.45 | 58.04 | < 0.001 |
|  | Population | 5.040 | 1 | 6.604 | 7.789 | 0.029 |
|  | Treatm. x Pop. | 0.648 | 1 | 60.01 | 1.002 | 0.321 |
| Fruit set | Treatment | 1.457 | 1 | 61.00 | 9.476 | **0.003** |
|  | Population | 2.405 | 1 | 61.00 | 15.64 | **< 0.001** |
|  | Treatm. x Pop. | 0.112 | 1 | 61.00 | 0.725 | 0.398 |
| Ovules per flower | Treatment | 0.020 | 1 | 12.21 | 0.524 | 0.483 |
|  | Population | 0.050 | 1 | 17.02 | 1.282 | 0.273 |
|  | Treatm. x Pop. | 0.007 | 1 | 12.55 | 0.183 | 0.677 |
| Seed set | Treatment | 0.056 | 1 | 48.29 | 0.267 | 0.608 |
|  | Population | 0.001 | 1 | 27.29 | 0.009 | 0.924 |
|  | Treatm. x Pop. | 0.140 | 1 | 49.75 | 0.666 | 0.418 |
| Seeds production per plant | Treatment | 11.42 | 1 | 50.00 | 21.48 | **< 0.001** |
|  | Population | 0.401 | 1 | 50.00 | 0.755 | 0.389 |
|  | Treatm. x Pop. | 0.012 | 1 | 50.00 | 0.022 | 0.883 |
| Pollen per anther | Treatment | 0.616 | 1 | 17.00 | 25.32 | **< 0.001** |
|  | Population | 0.004 | 1 | 17.00 | 0.158 | 0.696 |
|  | Treatm. x Pop. | 0.093 | 1 | 17.00 | 3.807 | 0.067 |
| Significant *P*-values were highlighted in bold. |  |  |  |  |  |  |

**Fig 4.**
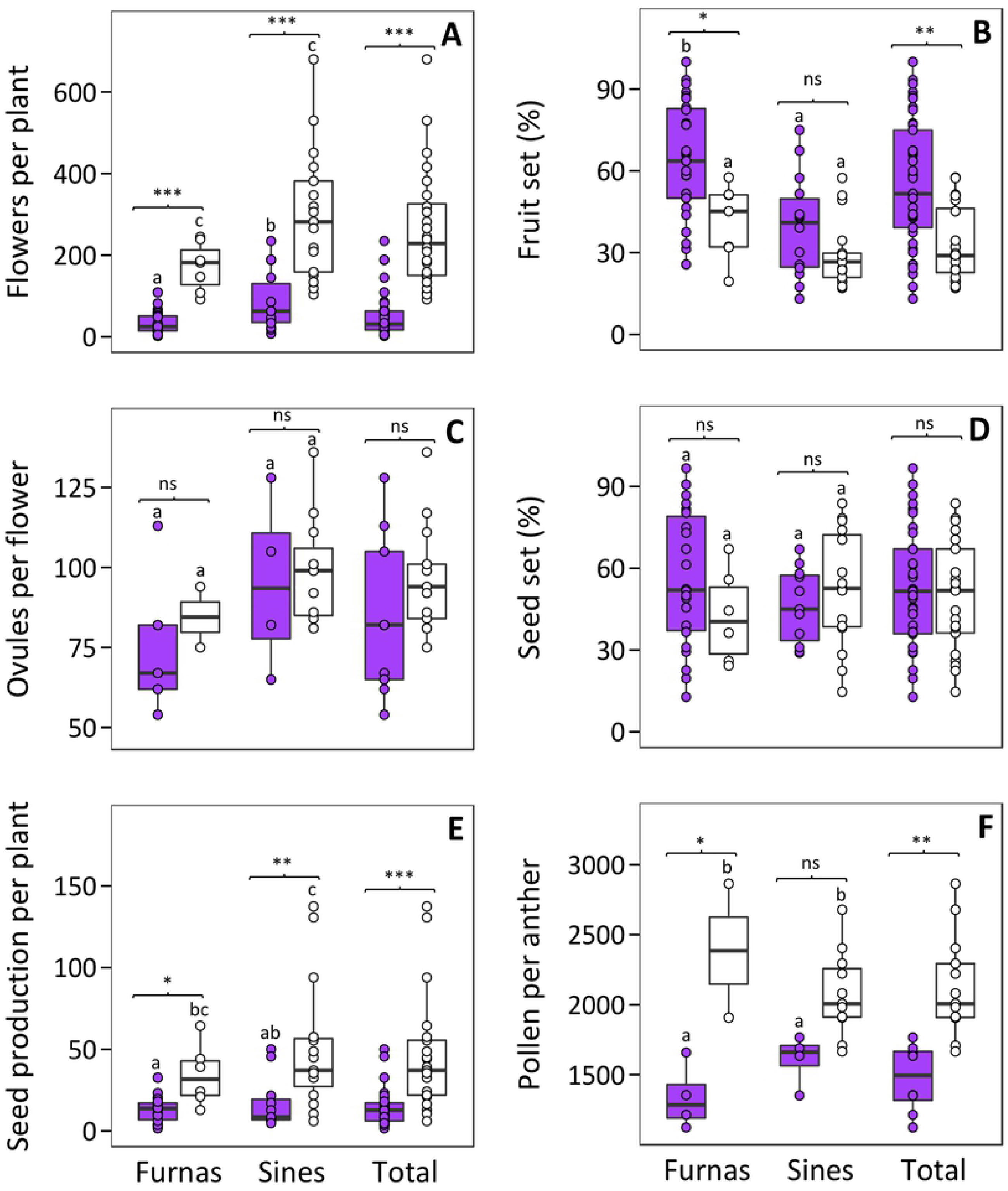
Boxplots representing the total flowers per plant (A), fruit set (B), ovules per flower (C), seed set (D), seed production per plant (E) and pollen per anther (F) in plants growing in the UV-present (purple boxes) and UV-exclusion (white boxes) treatments. Dots represent values for all estimations of plant reproductive performance. The central line displays the median, the bottom and top of the box are the first and third quartiles, and dots represent sample values. Letters displays are used to show results of multiple comparisons between populations. Within each population, pairwise comparisons between light treatments using Bonferroni adjustment are showed. ns, not significant; *, *P* < 0.05; **, *P* < 0.01; ***, *P* < 0.001.

When assessing the relationship between flavonoid production and male and female reproductive outputs, we did not found any significant correlations at Bonferroni-corrected level (α = 0.05/6 = 0.008) (S2 Table).

## Discussion

### Effects of UV radiation on flavonoid production

Exposure to UV radiation led to a generalized increase in the anthocyanin and flavone concentrations in *S. littorea*, suggesting that increasing flavonoid concentrations is part of this plant’s response to UV stress. In this species, all anthocyanins in aboveground tissues are cyanidin derivatives, whereas the most abundant flavones are apigenin derivatives (isovitexin) in petals and luteolin derivatives (isoorientin) in photosynthetic tissues [38]. Isoorientins are dihydroxy B-ring-substituted flavonoids, which it is known to have higher antioxidant properties [14,51–53]. In this regard, the UV-induced accumulation of efficient antioxidant flavonoids has been previously described in plants. For example, the flavonoid composition in leaves of the white clover (*Trifolium repens*) is affected by solar radiation, as the concentration of quercetin increases under UV-B stress rather than those of less effective antioxidants such as monohydroxylated kaempferol glycosides [54]. In addition, most anthocyanins and flavones in *S. littorea* are acylated [38], which it is known to enhance the flavonoid absorption in the UV-A and UV-B wavelength [28,55,56]. In *Cistus salvifolius*, for example, the occurrence of acylated flavonoids in trichomes ameliorates stress from excess UV radiation [57], whereas leaves of purple basil (*Ocimum basilicum*) accumulate coumaroyl anthocyanins that are more responsive to quenching sunlight irradiance, mainly in the UV-B wavelength, than non-acylated anthocyanins [27]. Although we did not obtain direct evidences of variations of the flavonoid-mediate quenching of UV-induced free radicals in *S. littorea*, its flavonoid profile, including acylated and efficient antioxidant flavonoids, suggests that this species show a robust biochemical toolkit that may protect itself from the oxidative stress caused by UV radiation.

Despite differences in flavonoid production caused by UV radiation, plants not exposed to UV light accumulated important amounts of anthocyanins and flavones in most aboveground tissues. This may be due to the incidence of high levels of PAR on these plants. Several studies indicate that UV irradiance is not a prerequisite for flavonoid biosynthesis. For example, the concentration of UV-absorbing quercetin in *Brassica oleracea* increases in line with PAR levels [58]. In *Arabidopsis thaliana*, PAR-only exposure contributes to the formation of a base amount of quercetin, providing a basic photoprotection that is further increased by long term exposure to UV-A/B radiation [59]. Similarly, high levels of PAR might lead to the formation of a base amount of anthocyanins and flavones in *S. littorea*, whose concentrations could be optimized and increased when plants are exposed to UV radiation. However, given that anthocyanins and flavones perform a plethora of protective functions against many biotic and abiotic factors [28,60,61], they could be performing non-photoprotective functions. For example, isoorientins and isovitexins found in flax (*Linum usitatissimum*) enhance the resistance to fungal infections [62], and petal isovitexins of *Silene latifolia* help regulate vacuole homeostasis in epidermal cells, preventing petals from wilting [63]. In addition, flavones are produced constitutively in aboveground tissues of *S. littorea* when plants grow in low light levels conditions [40]. Thus, although anthocyanins and flavones are probably conferring protection against high levels of PAR, we cannot rule out that the selective pressures of other biotic and abiotic agents could explain the constitutive accumulation of flavonoids found in plants not exposed to UV stress.

The increase of flavonoid accumulation in response to UV radiation was not homogeneous across tissues: petals respond to UV by increasing anthocyanins, calyces and leaves respond by increasing flavones, and stems through both anthocyanin and flavones. This result is not surprising because the biosynthesis of flavonoids is tissue-specific regulated [64]. The depletion of anthocyanins in petals en el UV-absent treatment is translated in a change color intensity [65], which may be differentially perceived by the insect pollinators [66]. On the other hand, calyces of plants from both UV-exclusion and UV-present treatments of Sines population showed higher levels of flavones than those of Furnas in each treatment. This difference may reflect a local adaptation of flavones biosynthesis to the higher UV radiation of the Sines population compared with Furnas [39]. However, further studies are necessary to assess whether flavonoid biosynthesis in *S. littorea* shows signals of local adaptation across the UV radiation gradient in the distribution area.

### Effects of UV radiation on photosynthetic performance

*Silene littorea* showed a higher decline of the quantum efficiency of PSII when plants were exposed to UV stress, especially in leaves. Many authors have showed that the UV part of sunlight is potentially highly important in photoinhibition of PSII of leaves. For example, *Cucurbita pepo* leaves under UV stress exhibit a parallel decrease in photosynthetic activity [42]. In addition, Albert et al. [67] showed that the PSII performance and net photosynthesis of *Salix arctica*, is negatively affected by the ambient solar UV-B radiation. Given that *S. littorea* was more susceptible to photoinhibition when it is exposed to UV stress, our findings add evidences that the ambient solar UV radiation is a significant stress factor for the photosynthetic activity of plants.

Despite the negative effects of UV stress on the photosynthetic activity in *S. littorea*, this species seems to have an optimal light-stress recovery system and does not incur in chronic photoinhibition, expressed as *Fv/Fm* values within the range of healthy plants after relaxation of photoinactivation. In plants, when incident light surpasses the energy assimilated by the photosynthetic apparatus, the excessive energy causes photoinhibition and the formation of ROS, which results in photo-oxidative damage and an eventual decline in photosynthetic activity [15,28,68]. The photoprotection mechanism of plants involves a variety of defense agents against light-induced ROS, including the synthesis of antioxidant anthocyanins and flavonoids [12,13]. In this regard, dihydroxy B-ring-substituted flavonoids located in the chloroplasts help antioxidant enzymes to reduce light-induced ROS and those diffusing out of the chloroplast are scavenged by vacuolar flavonoids [15]. In addition, leaves accumulating anthocyanins incur in less photoinhibition after a saturating light stress as compared with green leaves [27,69]. We hypothesized that flavonoids (both anthocyanins and flavones) of *S. littorea* may contribute to photoprotection to thrive in habitats with highly solar radiation such as coastal foredunes along the Iberian Peninsula [39]. Thus, flavonoid biosynthesis may be of particular benefit to *S. littorea* to prevent photoinhibition in this light-stressed habitat, as it was found in *Silene germana* [70].

### Effects of UV radiation on reproductive output

Plants exposed to UV produced approximately three times less total number of seeds per plant than those shielded from UV, driven primarily by a decrease in total flower production. In a previous study, we found that flower production in *S. littorea* increases as a consequence of high natural sunlight levels [71], but exposure to sunlight also entails the exposure to harmful UV wavelengths. Here, we demonstrated that the absence of these harmful effects in the UV-exclusion treatment allows the absorption of PAR and enhances flower production. Although many studies often report enhanced flowering when plants were exposed to supplemental UV radiation (e.g. [72,73]), other studies have reported the opposite effect. For example, Feldheim and Conner [74] reported that supplemental UV-B radiation was generally detrimental to flowering in *Brassica nigra* and *B. rapa*, while plants from a lowland population of *Silene vulgaris* increase their flower production in the absence of UV-B [75].

Additionally, we found that the proportion of flowers yielding fruits was nearly double in plants under UV stress. Even though other studies have reported increasing fecundity in plants exposed to moderate supplemental UV radiation [76], we suggest that significant differences in fruit set between light treatments could be influenced by the resources allocated to the high flower production of plants growing in the absence of UV stress. In addition, pollinators can become saturated at high flower densities [77,78], resulting in a decrease of per-flower visitation. Thus, despite the fact that absolute fruit production was almost four times higher in plants shielded from UV light, the elevated number of flowers of these plants not visited by pollinator may have led to a reduced fruit set. Experimental plants were fully accessible by pollinators around the study area (mostly hymenopterans), thus we can rule out that any architectural effect of the experiment might difficult pollinator visits.

Pollen production decreased in plants exposed to UV light is consistent with results in other species [36,37,79]. Since male gametes of plants are encased in pollen grains, decreasing pollen production is expected to have an adverse impact on male fitness of *S. littorea*. Conversely, ovule production was similar in plants from both light treatments. Ovules occur in ovaries, which are well protected against UV stress due to their accumulation of UV-absorbing compounds that attenuate UV radiation [7,33]. In *S. littorea*, upper anthers occur slightly beyond the corolla opening at anthesis and are more exposed to UV radiation, whereas carpophore is embedded in the calyx. Thus, ovule production is less likely to be compromised by solar radiation since ovules are protected from UV radiation by several layers of tissue.

## Conclusions

UV radiation incurred a trade-off between flavonoid protection and reproduction in *S. littorea*. We propose that flavonoid production was activated as a defense mechanism against UV radiation, presumably because of their antioxidant nature, which may prevent the chronic photoinhibition and promote a rapid photosynthetic recovery. Conversely, exposure to UV radiation negatively affected flower and pollen production in this species. This balancing between protection and reproduction may be beneficial to successfully survive in exposed coastal foredunes. Thus, the allocation of metabolic resources may provide an efficient photoprotective toolkit and, at the same time, guarantee the reproduction of this species in Mediterranean climates subjected to strong UV radiation.

## Acknowledgments

The authors thank to A. Gallardo for their laboratory assistance and A. Cardoso for her help at the greenhouse. We particularly thank the anonymous referees for their helpful comments. This work was supported by the Spanish Government MINECO projects (CGL2012-37646 and CGL2015-63827-P) and a Predoctoral Training Program grant to JCV (BES-2013–062610).

## Supporting information

**S1 Table. Number of plants for each maternal genotype, population and treatment (UV-present and UV-exclusion treatments).** The number of plants sampled for anthocyanin and flavone concentration is indicated in parentheses.

**S2 Table. Pearson correlation coefficients for the comparison between flavonoid production (anthocyanins and flavones) in each plant tissue and reproductive outputs of *S. littorea*.**

